# High-throughput Experimental Validation of Synonymous Mutation Neutrality in the Human Genome

**DOI:** 10.1101/2025.08.30.673250

**Authors:** Yiyun Rao, Ivan Sokirniy, Edward O’Brien, Justin Pritchard

**Author notes:** Corresponding Author: Edward O’Brien, Justin Pritchard.

## Abstract

The assumption that synonymous mutations are fitness-neutral is central to many foundational results in the fields of genetics, genomics, evolutionary biology, and medicine. However, recent results suggest synonymous mutations have pervasive and strong fitness effects. These vigorously debated studies in non-human model systems have even suggested that the proportion of synonymous mutations and their fitness effect sizes are similar to non-synonymous mutations. To systematically test synonymous neutrality across the human genome we utilize recent advances in base editing to probe the fitness effects of 8,558 potential synonymous mutations in 128 highly essential genes. Combined with novel base editing controls and extensive individual sgRNA guide-level validation experiments that quantify false positive rates, we find that synonymous mutations rarely have fitness effects on growth, occurring 165-fold (95% CI, 35-2929 fold) less frequently than non-synonymous fitness-altering mutations. Thus, despite a decade of high-profile controversies, the neutrality assumption for spontaneous synonymous mutations is valid.

## Main

Synonymous mutations are nucleotide changes in codons that do not alter the encoded amino acid sequence. The assumption that synonymous mutations are largely invisible to natural selection is a cornerstone of modern protein-based phylogenetic models^1,2^. It is embedded in landmark genomic annotation efforts^3,4^, been incorporated into foundational phylogenetic methods^4^ that have shaped our understanding of the functional elements in the genome and form the basis for identifying and cataloging cancer driver genes^6,7,8^ that serve as a reference for biomedical research and therapeutics^7^ in cancer genomics.

In the past 15 years, a string of high-profile and hotly debated studies have provided evidence that challenges the neutrality assumption. Research from a range of model organisms has suggested that at least some synonymous mutations in some genes might have a distribution of fitness effects that is similar to non-synonymous mutations^8–10^. Evidence has been found for both deleterious^9,11^ and adaptive^12,13^ fitness changes. In 2022, a controversy arose from a study claiming that 70% of synonymous mutations in 22 essential yeast genes are non-neutral, a proportion comparable to nonsynonymous mutations^10^. This surprising finding was challenged due to methodological concerns^14^. In response, the authors published a follow-up study with more appropriately controlled experimental data^11^ and found they had overestimated the magnitude of the fitness effects but defended their conclusion that the majority of synonymous mutations are non-neutral. This collection of results is provocative because violating the assumption of synonymous neutrality would call much of evolutionary biology, bioinformatics, and disease genetics into question. Because of the stakes, and the literature’s use of non-human model systems, we believe a direct experimental interrogation of the fitness effects of select synonymous mutations in human cells within the native genomic context is critical to either support or reject the assumptions of fitness neutrality for synonymous mutations.

### Framework for Isolating Synonymous Variant Effects in Pooled Screens

While there are no systematic assessments of synonymous variant function in human cells, several recent large-scale genome editing studies report that synonymous mutations exhibit extensive non-negligible fitness effects. For example, 21%^15^ of synonymous mutations in the BRCA1 gene appear to be loss-of-function variants as determined via a deep mutagenesis assay. In contrast, a tiling base-editing assay estimated this same proportion to be 1.2%^16^. Similarly, base editing screens targeting ‘core’ fitness^17,18^ genes reported that between 5.2%^19^ (in 7 genes) and 11.5%^20^ (in 10 genes) of sgRNAs directed at synonymous sites created fitness defects that caused the frequency of synonymous editing sgRNAs to deplete in the population. This wide range of reported values (1.2%–21%) likely stems from the fact that these studies’ primary focus was on characterizing non-synonymous mutation effects. It is therefore possible that the observed significant synonymous sgRNA hits are less reliable than the non-synonymous results because the studies did not explicitly attempt to provide controls for these synonymous hits or validate them in follow-up assays.

In designing a new base editing study to directly interrogate the fitness consequences of specific synonymous mutations at the endogenous genomic locus in human cells, we identified three potential sources of false positives (Figure 1A). First, unintended non-synonymous edits due to bystander mutations beyond a base editor’s core editing window. Most base editing studies report only the highest probability 5-nt ‘core’ editing window within the sgRNA sequence. This region, typically spanning nucleotide positions 4–8 relative to the protospacer adjacent motif (PAM), exhibits the highest deaminase activity and editing efficiency^25,26^, however, editing activity occurs outside this core window in at least 20% of cases ^21^. This means that many guides that are labeled as “synonymous” guides in existing studies make only synonymous mutations in the smaller 5-nt high probability editing window, but, have a significant potential to make non-synonymous mutations just outside this window. Second, Cas9-independent off-target effects caused by base editors acting on single-stranded DNA could produce mutations randomly in the genome^22,23^. Even though the rate of such an effect could be estimated by safe harbor targeting guides^20^, the resulting false positive effect would still appear in a pooled screen. Finally, there are potentially overlooked editing independent effects from Cas9 nickase activity disrupting essential gene expression^24^. Cas9, for example, can repress transcription by obstructing initiation or elongation when sgRNAs target regions near promoters ^24,25^. The extent of this repression is influenced by factors such as the sgRNA’s position relative to the promoter, guide composition, chromatin state, and the concentration of the Cas9/sgRNA complex^26^.

**Figure 1.**
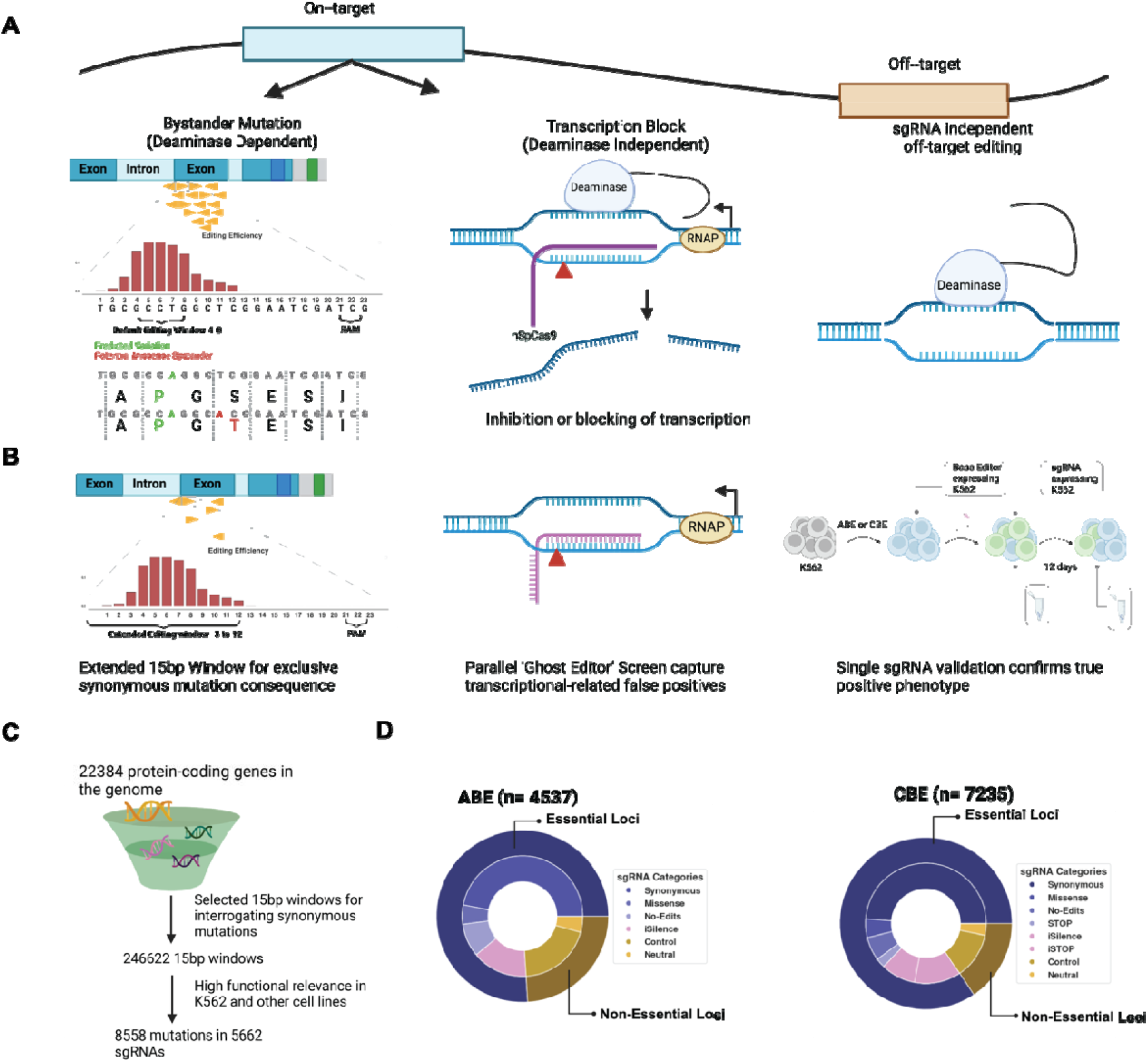
Minimizing False Positives in Base Editor Screens of Synonymous Mutations (A) Key sources of false positives in pooled sgRNA screens: 1. Bystander Mutations (Deaminase-Dependent): Base editing can introduce bystander mutations within the default core editing window (typically 4-8 bp), resulting in unintended missense mutations. This confounds the intended synonymous mutations, leading to false positive phenotypes. 2. Transcriptional Block (Deaminase-Independent): The binding of nCas9 to single-stranded DNA can inhibit transcription initiation or elongation, potentially reducing gene expression. This deaminase-independent effect can mimic gene knockdown, producing false positive depletion phenotypes.3. Off-Target Editing: sgRNAs may bind to off-target sites in the genome, leading to unintended base editing at functional loci. This confounding variation can falsely suggest gene essentiality or an altered phenotype (B) Strategies to isolate the effects of synonymous mutations:1. Extended Editing Window: By expanding the base editing window from the default 5 bp to 15 bp, sgRNA designs can reduce the likelihood of bystander missense mutations, increasing the chance of inducing pure synonymous changes.2. ‘Ghost’ Editor Control: A parallel screen using a deaminase-null base editor (referred to as the ‘Ghost Editor’) should be performed with the same sgRNA library. This controls for transcriptional block effects, enabling the identification of transcription-related false positives.3. Single sgRNA Validation: Hits from pooled screens should undergo validation with individual sgRNAs to confirm true positive phenotypes and distinguish them from false positives arising from off-target or bystander effects. (C) Diagram of selecting sgRNA for an unbiased screen. A total of 128 target genes were selected for screening. (D) Composition of ABE and CBE sgRNA libraries. The ABE library consists of 4537 sgRNAs, including synonymous sgRNAs, missense sgRNAs, sgRNAs designed to make no edits in the same essential gene, and positive controls such as sgRNAs targeting start codons (iSilence) in essential loci. In non-essential loci, neutral sgRNAs target K562-neutral genes, and control sgRNAs include non-targeting sgRNAs. The CBE library consists of 7235 sgRNAs. In addition to the categories in the ABE library, the CBE library includes sgRNAs that induce stop codons in essential genes (iSTOP) and an iSTOP positive control library.

To address these issues, we implemented a four-pronged validation framework not used in previous studies (Figure 1B): (1) we increased the base editing window to 15 bp to ensure that our “synonymous” sgRNAs produced exclusively synonymous mutations across the entire editing window, not just the 80% of edits in a 5 bp window; (2) We used a Ghost Editor Control, which is a deaminase-null base editor, thus any sgRNA hits from the library in this deaminase deficient screen are editing-independent false positives because no mutations are made. (3) We carried out independent infections of the library to control for Cas9-independent editing effects. And (4) we validated significant hits from the pooled screen with single sgRNA assays followed by direct sequencing to detect any off-target edits.

To establish a large sgRNA library to systematically investigate the fitness effects of synonymous mutations across the human genome, we aimed to maximize the number of “synonymous mutation only” sgRNAs within highly essential genes across multiple cell lines using the efficient and specific NG-PAM base editors, ABE8e^27^ and CBEd^28,29^. Starting with 22,384 protein-coding genes in the genome, we applied a data-driven approach to identify regions of interest. Specifically, we identified all 15 bp windows within coding regions that were devoid of the possibility of non-synonymous edits, thereby focusing on areas where synonymous mutations could be interrogated for their potential functional impacts. This resulted in 246,622 windows that we could target (115,401 targeted by the ABE base editor and 131,221 targeted by the CBE editor). Next, we restricted this set to 128 genes that were highly essential in both of our cell models by DepMap^17^ gene effect score and broadly essential across diverse cellular contexts and created a sgRNA library that can target 5,662 windows (2,134 in ABE and 3,528 in CBE) with the potential to introduce 8,558 putative synonymous mutations (2,964 targeted by ABE and 5,594 targeted by CBE; Supplementary Table 1-8). There are more putative synonymous mutations than sgRNAs because a single guide has the potential to create multiple mutations within the target window (Figure 1C).

To ensure a robust experimental design, we incorporated rigorous controls into the sgRNA library. Negative controls included sgRNAs with no predicted edits within the exact same gene (*n*=407 in ABE, *n*=458 in CBE), non-targeting sgRNAs (*n*=613 in ABE and CBE), sgRNAs targeting safe harbor sites (*n*=282 in ABE and CBE) and neutral sgRNAs targeting non-essential genes (*n*=196 in ABE, *n*=231 in CBE). Positive controls included sgRNAs that create missense mutations in the same essential genes (*n*=232 in ABE, *n*=367 in CBE), sgRNAs that create premature stop codons in the same essential genes (*n*=181 in CBE), sgRNAs designed to disrupt start codons (known as iSilence, *n*=673)^30^ in both the ABE and CBE libraries, and previously published iSTOP sgRNAs (*n*=902)^31^ in the CBE library. The final library contains 4,537 and 7,235 sgRNAs, respectively, for the ABE and CBE libraries, which are composed of controls, synonymous and missense mutation causing sgRNAs (Figure 1D, see Supplementary Note for Library Composition, Supplementary Table 3 and 4).

To mitigate base editor-induced off-target effects and cell line-specific effects, we independently transduced three replicates of K562 and Jurkat cells, with the sgRNA library on a separate vector (denoted D0). Following the library introduction, replicates were again independently infected with ABE8e, CBEd, or the deaminase-null Ghost Editor. After infection and selection, the first endpoint was day 12 (D12) later for both ABE and CBE screens in K562 and Jurkat cells (Supplementary Figure 1). Given the lower editing efficiency of cytosine base editors compared to adenine base editors ^32^, the CBE screen duration in K562 was extended by an additional 14 days (D26) to allow for the potential to identify more CBE hits. Cells were then harvested for next-generation sequencing (NGS) to assess sgRNA representation at the baseline and Day 12 or Day 26 endpoint. Differential read counts were analyzed using DESeq2

### Functional synonymous variants are extremely rare in human cells

Positive control libraries (iSilence), as expected, exhibited a high concordance of normalized log2 fold changes (L2FC) across biological replicates (Supplementary Figure 2,3,4), with Pearson’s r = 0.87 in Jurkat and 0.83 in K562 for ABE8e. For CBEd, Pearson’s r was 0.64 in K562 and 0.79 in Jurkat at the first time point, confirming consistency across replicates. We observed iSilence hit rates, defined as the number of significant (adjusted p-value < 0.05) sgRNAs out of total sgRNAs screened within the set, of 30.6% in Jurkat and 32.3% in K562 for ABE8e Negative control sgRNAs set in both screens showed minimal hit rates, ranging from 0% to 4%. (Supplementary Figure 2,3,4). These results are consistent with the prior literature and indicate a successful screen^20^.

**Figure 2.**
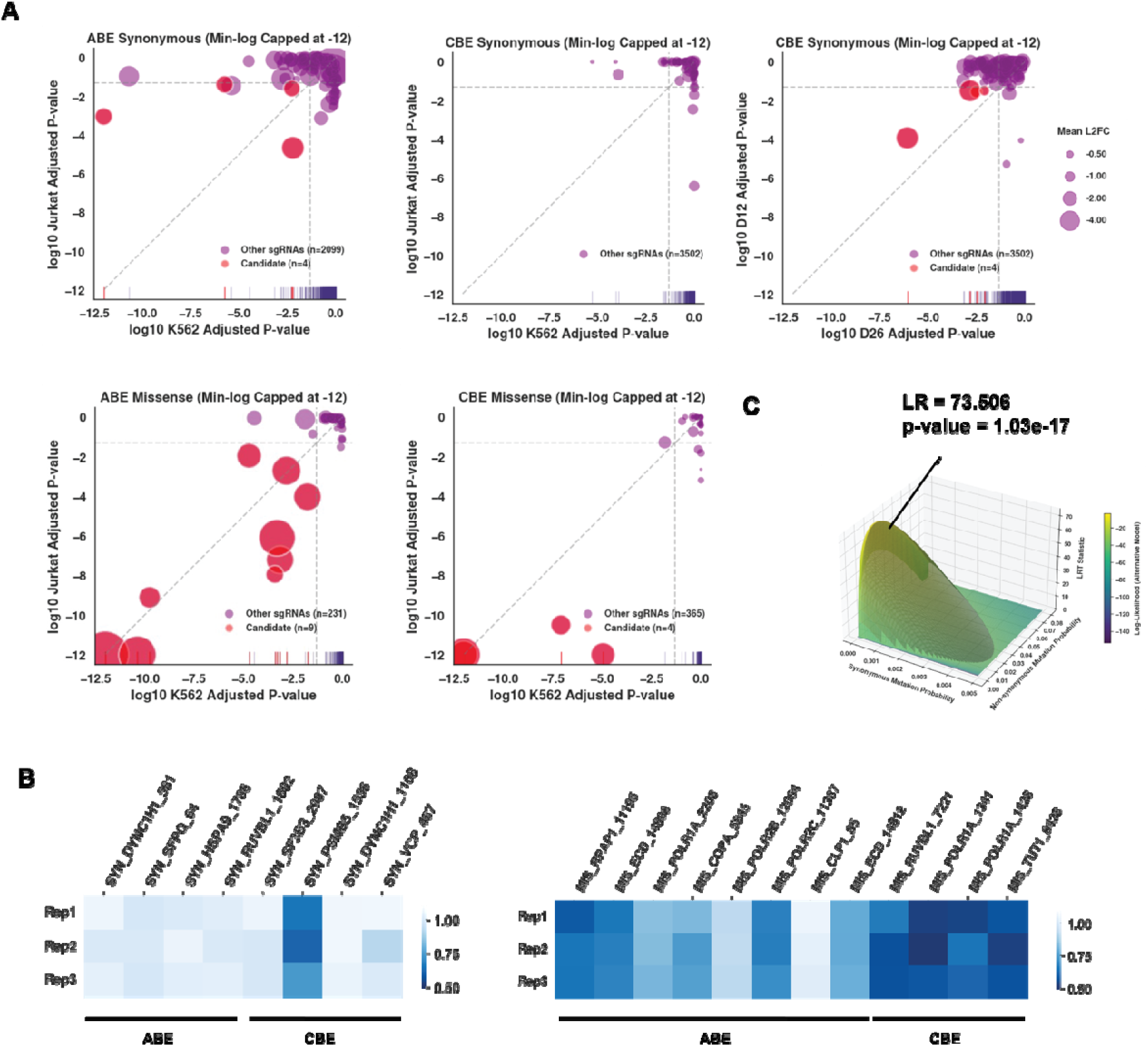
Experimental Validation of Synonymous sgRNA hits (A) The x-and y-axes represent the –log10-transformed adjusted p-values of the same sgRNA library in different conditions: (1) K562 vs. Jurkat for ABE synonymous and missense sgRNA, and CBE missense candidates, and (2) Day 26 vs. Day 12 for CBE synonymous candidates, as there were no overlapping candidates between K562 and Jurkat. Each dot represents a sgRNA, with red dots highlighting qualified candidate sgRNAs. Dot size is proportional to the average log2 fold change in the respective conditions. For visualization purposes, the –log10-transformed p-values are capped at 12. (B) Heatmap of GFP+% cell population for tested sgRNA for all biological replicates. The color shows GFP+% of Day 17 normalized by Day 5. A deeper blue indicates a larger depletion. (C) The likelihood ratio surfaces, where the x-axis and y-axis represent probabilities of synonymous and non-synonymous mutations having fitness effects, respectively. The z-axis represents the likelihood ratio, and the color gradient encodes the combined likelihoods.

The strong performance of the screen in our positive and negative control guide sets gave us confidence that we could analyze the non-synonymous and the synonymous hits from the screen. To do this, we looked for guides that were significantly enriched or depleted on day 12 and/or day 26. After identifying hits in both K562 and Jurkat cells, we filtered out for guides that were also hit in our ghost editor screen (Supplementary Note, Supplementary Figure 5) as they are likely false positives. We ended up with a sgRNA hit rate of 2.17% (13/599) and 0.141% (8/5,662), respectively, in non-synonymous and synonymous mutation-inducing guides (Figure 2A, Supplementary Note).

As the final validation step in our approach, we performed single sgRNA competition assays (mutant versus wild type) across multiple time points on 20 of the most significant hits across both CBE and ABE screens, corresponding to 12 non-synonymous and 8 synonymous sgRNAs. To perform these experiments, we established K562 cell lines stably expressing base editors and GFP-tagged sgRNA vectors via lentiviral infection. This created a mixed population of GFP-negative (base editor only) and GFP-positive (base editor + sgRNA-expressing) cells. We monitored the proportion of GFP+ cells and the edited allele frequency over 12 days to assess depletion effects. While all sgRNAs produced the expected edits, only 1 out of 8 synonymous hits depleted in this validation assay (Figure 2B), indicating 7 were false positives in the pooled screen. For the synonymous hit that validated we also performed whole genome sequencing to identify off-target edits that might produce missense mutations. We identified 2 missense mutations in essential genes that occurred in all replicates, but these mutations did not deplete over time, suggesting that they occurred in a distinct subpopulation and did not have a fitness defect (Supplementary Note). This strongly suggests that the single validated synonymous edit is a true positive. In contrast, 11 of the 12 missense sgRNAs across both the ABE and CBE libraries exhibited significant depletion by GFP (Supplementary Figure 6,7) and NGS (Supplementary Note), and all missense hits introduced the target mutation according to whole genome sequencing. Thus, our validation assays revealed a 91% true positive rate (11/12) for missense sgRNAs that were scored as hits in our pooled screen, but only 12.5% of the synonymous sgRNAs that were scored as hits in our pooled screen were true positives (1/8).

### Estimation of the relative probability of synonymous fitness effects

We next sought to estimate the relative probability that a synonymous mutation will have a fitness effect compared to a non-synonymous mutation. To calculate this, we use three values: 1) The number of mutations expected to be introduced by our sgRNA library. 2) The number of significantly different measurements, *i.e.*, ‘hits’, in the high throughput, pooled assay. 3) The number of validated mutations contributing to the phenotype measured in our low-throughput, validation assay. For our synonymous library, 8,558 potential synonymous mutations yielded 8 sgRNA hits in the pooled assay, with one mutation from a single guide confirmed to affect phenotype in the validation assay. This suggests that 0.0117% (1/8558) of our potential synonymous edits impact fitness. In contrast, our non-synonymous library tested 878 potential non-synonymous mutations of which 17 had validated fitness effects, suggesting that 1.93% of non-synonymous edits have fitness effects. This difference of 0.0117% versus 1.93% indicates that non-synonymous mutations are approximately 165-fold more likely to have a fitness effect than synonymous mutations. The 95% confidence interval estimated by the joint distribution of synonymous and non-synonymous mutation fractions is 35.02 to 2,929.

We next asked whether the distribution of fitness effects differs between synonymous and nonsynonymous mutations. To do this we applied a likelihood ratio test to compare the validated hit rates between synonymous and non-synonymous sites, given the null hypothesis that the distribution of fitness effects is the same between synonymous and non-synonymous mutations, (Figure 2C). Likelihood surface and test statistics supported a significant difference between these categories with a test statistic of 73.51 and a p-value of 1.03×10^-17^. Thus, we reject the null hypothesis that synonymous and nonsynonymous mutations have equal probabilities of conferring a fitness effect.

While the likelihood ratio suggests that synonymous and non-synonymous mutations do not have similar effects on fitness, as suggested elsewhere^34^, we also wanted to evaluate whether synonymous mutations tend to behave neutrally. To do this we compared the hit rate distributions between synonymous-targeting sgRNAs and multiple categories of negative control sgRNAs (non-targeting, safe harbor, and neutral-targeting). The interquartile range (IQR) of hit rates for negative control sgRNAs spans from 0.34% to 3.43%, with a median of 1.03%. Synonymous-targeting sgRNAs exhibited a similar median hit rate of 0.95%, with values ranging from 0.22% to 1.22%. Grouping negative control sgRNAs did not alter the observed similarity in distribution. When grouped, negative control sgRNAs showed an interquartile range of 0.26% to 2.2%, which remains comparable to that of synonymous-targeting sgRNAs (Supplementary Table 12). Therefore, our result indicates that guides that create synonymous mutations behave similarly to guides that are negative controls. This suggests that synonymous mutations are predominantly neutral in human cells.

To the best of our knowledge, these results are the first widespread and direct confirmation that non-synonymous mutations made at endogenous human genome loci are significantly more likely to have a fitness impact than synonymous mutations.

## Discussion

Our study employed a novel four-pronged approach to estimate the fitness effects of synonymous mutations in essential genes within their endogenous genomic context using base editing. We designed our library with stringent criteria, selecting 15-bp windows in essential genes that permit only synonymous edits, thereby eliminating the possibility of confounding non-synonymous mutations. We included a novel deaminase-deficient editing control that identified and then filtered out the base edits that score in our screens in the absence of a functional editor, and we validated depletion phenotypes and base edits in independent validation assays that were not included in several other studies^19,20^. The similar hit rates observed for synonymous and non-synonymous mutations in those studies may be attributable to the absence of such controls and lack of follow-up validation. Indeed, our validation assay revealed a high false-positive rate (91%) among the limited significant synonymous hits from the pooled screen.

Collectively, our results demonstrate that single synonymous mutations in the human genome are 165-fold (95% CI, 35-2929 fold) less likely than non-synonymous mutations to impact fitness. This contrasts with claims based on non-human model organisms that the functional effects of synonymous mutations are similar to non-synonymous mutations^10,11^. The high performance of positive controls in our study that induce strong fitness effects by eliminating start codons, creating stop codons, and introducing missense mutations gives us confidence in the quality of our base editing measurements. Indeed, comparing the effectiveness of our positive controls to the small number of significant synonymous hits in our carefully designed synonymous library creates a striking observation of the rarity of synonymous fitness effects.^11,12^

Our study has focused on measuring differences in cellular fitness conferred by individual or a small number of mutations made by single sgRNAs, not the influence of epistatic interactions between mutations. Indeed, when large swathes of codons within a gene are simultaneously, synonymously mutated significant phenotypic changes have been observed^35,36^. Moreover, our study did not probe changes in protein co-translational folding^37,38^, mRNA expression^39^, stability and degradation^40,41^, or other changes^42^ associated with codon optimization. If such changes in molecular phenotype are widespread, our observed rarity of fitness effects from synonymous mutations suggests that these molecular perturbations are effectively buffered by multiple layers of homeostasis mechanisms.

Taken together, our work constitutes the first large-scale experimental test of synonymous neutrality in the human genome, and the results confirm that the assumption of synonymous fitness neutrality, relative to other mutation types, is a reasonable approximation in most cases. In light of recent claims from other studies, this result reassures us that many of the foundational results and widespread analysis methodologies in genetics, genomics, evolutionary biology, and medicine are correct.

## Method

### Dataset

The K562 gene effect table, common essential gene list, common non-essential gene lists, and K562 transcript expression file were downloaded from DepMap 22Q2 (https://depmap.org/portal/data_page/?tab=allData). Canonical transcripts were retrieved from the MANE (Matched Annotation from NCBI and EBI) Project (https://www.ncbi.nlm.nih.gov/refseq/MANE/). Gene and transcript annotation were retrieved from GenCODE release 38 (https://www.gencodegenes.org/human/release_38.html). Protein-coding gene names across the whole genome were retrieved from BioMart(https://useast.ensembl.org/info/data/biomart/index.html)

### Essential Gene Selection

For essential genes, we selected genes in K562 that are < 0.01 quantile of the gene effect table and overlap with the common essential genes, resulting in 128 candidate genes (Supplementary Table 1). For non-essential genes, we selected genes with a gene effect score between -0.01 and 0.01, intersecting with the common non-essential genes, resulting in 41 non-essential genes (Supplementary Table 2).

### sgRNA Library Design

The selected essential gene list was used to run CHOPCHOP locally with a setup based on GRCh38 Genome Assembly. All possible sgRNAs with NG PAM in the coding region, with padding size 0, were returned. An updated setup for the Mac M1 Pro chip and the script can be found on GitHub (https://github.com/ryy1221/synSg).

To filter the unqualified sgRNAs, first, sgRNAs that can bind off-target sequences with fewer than two mismatches were filtered out. Then, sgRNAs overlapping with an intron-exon boundary were filtered out. For the MANE canonical transcript of each gene, the 15bp base editing window (defined as -3 to +12 positions of the protospacer binding sequence, where 1 is the most PAM-distal position, and positions 21-23 are the PAM) was identified, and all possible edits within the window were predicted. To avoid confounding effects from canonical splicing, all sgRNAs targeting positions within ±2 base pairs of the intron-exon junction were excluded. The amino-acid alteration consequences of all possible edits were then predicted based on the hg38 annotation file.

For each gene (both essential and non-essential), the categories of sgRNAs were defined as:

1. **Synonymous:** All possible edits within the window generate synonymous mutations.
2. **Missense:** >=1 edits within the window generate missense mutations.
3. **Stop:** >=1 edits within the window generate nonsense mutations.
4. **Empty Window:** No edits were predicted in the target region.

We order the mutation types as Nonsense > Missense > Silent. Guides containing multiple mutation types were categorized by the most severe mutation type. All qualified synonymous sgRNAs were kept in the library.

To maintain a stringent definition of synonymous mutations, our analysis focused on synonymous mutations that do not overlap canonical splice sites. Specifically, we excluded positions within ±2 nucleotides of intron-exon junctions harboring the conserved AG-GU splice site motif. A minority of sgRNAs (4.35% in the ABE library (93/2,134) and 5.64% in the CBE library (199/3,528)) target sites within 8 base pairs of exon-intron boundaries. Our approach does not eliminate potential effects mediated by exonic splicing regulatory elements, such as enhancers or silencers (ESEs or ESSs).

Base editors have an “editing window”, a small range of nucleotides surrounding the target site where editing efficiency is maximized. When a sgRNA targets multiple codons, especially if these codons are spaced apart, it’s unlikely that all will fall within these high-efficiency windows. Therefore, for sgRNAs predicted to cause three or more mutations, we conservatively capped the total countable mutations at two for estimating the putative number of mutations introduced. This is a conservative estimation of mutation numbers given the validated NGS mutations.

### sgRNA and base editor vectors

The sgRNA libraries were cloned into the lentiCRISPR-v2-HygR-mCherry vector (Addgene #104991). For single sgRNA validation, individual sgRNAs were cloned into the lenti-ccdB-Dang-Hyg-sgGFP vector (Addgene #235047). The base editors used were ABE8e-SpG-Puro (Addgene #235044) for the ABE screen and CBEd-SpG-UGI-Puro (Addgene #235045) for the CBE screen. The constructed deaminase-null control, Ghost Editor, was nSpG-UGI-Puro (Addgene #235046).

### sgRNA Library Cloning

The final oligo sequences were attached with BsmBI sites and PCR primers following a modified version of the Doench Golden Gate cloning protocol. Every two sgRNAs were combined into one oligo, following the example: ForwardPrimerTCTCACACC-sgRNA1-TCCGTTTCGAGACGCGTCTCACACC-reverse complement of sgRNA2-AGTTTCGAGA-ReversePrimer An extra G was added to the start of the sgRNA if the sgRNA starts with C or T for enzyme digestion efficiency.

Primers for the ABE library:

- Forward Primer: GGGATCACTTACTACTTCCG
- Reverse Primer: GGTACCCTATCACTTAACCG

Primers for the CBE library:

- Forward Primer: CACTAGAGTAATCGCTACCG
- Reverse Primer: GGGCTACTAGACATAACTCG

Primers for the Dual iSilence library:

- Forward Primer: CCCTCTAGTCTTTCCAATCG
- Reverse Primer: GCCTGATATCACTCCTATCG

The libraries were synthesized by TwistBioscience.

The sgRNAs were amplified using KOD Hot Start DNA Polymerase (Sigma, 71086-3) and ligated (NEB, R0734S) to predigested lentiCRISPR-v2-HygR-mCherry vector via Golden Gate cloning.

### sgRNA library Transformation

One day before electroporation, inoculate 10-beta competent cells (NEB, C3019H) in 200ml media(20g Tryptone/L, 10g Yeast Extract/L). After the cell reached OD ∼=0.,5-0.6, cells were collected and washed in 10% glycerol 3 times (2500g, 15min). Cells were resuspended in 10% glycerol after 3 washes. Per electroporation reaction, 10uL ligation product was electroporated into 400uL prepared 10-beta electrocompetent cells using Eppendorf Eporator (P2, 2500V). The electroporated cells were recovered at 37C with 1ml SOC for an hour and transferred to 200LB media at 30°C for 16 hours with 100 μg/ml carbenicillin. Electroporation efficiency was determined by plating 100X and 1000X dilutions on carbenicillin plates after recovery to ensure a library coverage of at least 500X.

### sgRNA Library Transfection and Infection

24 hours before transfection, HEK293T cells were seeded in 10 cm dishes at a density of 0.5×10^6^ per ml in 10 ml of DMEM + 10% FBS. sgRNAs were transfected into HEK293T cells on day 1 with Gen2 lentiviral vector using calcium phosphate. Five 10 cm dishes of HEK293T cells were transfected with a total of 20 μg sgRNA library plasmid and 20 μg lentiviral v2 helper plasmids per plate. Media were changed 6 hours after transfection and again on the second day. The viral media were collected 2 days after transfection, and at least 14M K562 and Jurkat cells were infected with 1 ml viral media for 5M cells, ensuring infection efficiency Multiplicity of Infection (MOI) <1 and sgRNA coverage >500X. Five days after infection, % mCherry positive cells were monitored and selected in 200 μg/ml hygromycin for another 7 days. Concurrently, 20 μg ABE8e, CBEd, and nSpG plasmids were transfected into HEK293T cells with 20 μg lentiv2 vectors, respectively to generate base editor viral media.

The day after hygromycin selection ended was set as D0, and at least 15M cells were collected as the baseline. On the same day, sgRNA-infected and selected cells were infected with base editors. Five days after infection, the cells were selected with puromycin (1 mg/ml for Jurkat, 1.5 mg/ml for K562). For the ABE arm, both cell lines were selected for 7 days, and cells were collected on Day 12. For the CBE arm, cells were collected on Day 12. K562 CBE arms were cultured for another 14 days and collected again on Day 26.

### Library Preparation for Sequencing

At least 10M cells were harvested for each library in each condition. Genomic DNA was extracted using phenol-chloroform^43^ following cell lysis. For PCR amplification, staggered PCR was used. Detailed PCR primers can be found in the Supplementary Note. gDNA was divided into 100 μL reactions such that each well had at most 10 μg gDNA. PCR amplification was performed with Ex Taq (TaKaRa, RR001A), as described in previous protocol conditions and volumes^44^. PCR amplicons were extracted using gel extraction (OMEGA, D2500) and quantified by Qubit. Amplicons of sgRNA libraries were submitted to Novogene for PE150 sequencing. ^4445^

### Single sgRNA Validation Experiment and Sequencing

All candidate sgRNA sense and antisense oligos with BsmBI sites were ordered from IDT. Oligos were annealed and cloned into lentiCRISPR-v2-HygR-GFP Vector. K562 cells were cultured and infected with base editor plasmids used in the screen. Three days after infection, the infected cells underwent puromycin selection for base editor-containing K562 strains. Single sgRNA plasmids were then transfected with lentiviral packaging plasmids. The base editor-containing K562 cells were infected with single sgRNA plasmids. Five days after infection, the GFP-positive cells were tracked using flow cytometry for 12 days until Day 17.

At least 1M cells were collected at the start and end of the tracking period. gDNA was extracted using the NEB Monarch gDNA extraction kit(NEB, T1120S). Amplicons were amplified using Takara Bio Ex Taq polymerase, with primers that have standard Illumina adapters attached. The PCR products were then loaded on a gel and purified by gel extraction. Sequencing was performed by Genewiz Amplicon-EZ. To analyze genome editing outcomes, we used CRISPResso to process sequencing data from edited cell populations. Paired-end FASTQ files were provided as input, along with the reference amplicon sequence and an assigned amplicon name. Default parameters were used. The output directory contained quantifications of editing efficiency and mutation types, enabling downstream analysis of genome editing events. We applied filtering criteria to retain only sequences with at least 1% of mapped reads and a minimum of 100 read counts. Sequence modifications were identified by aligning observed sequences to reference amplicons and pinpointing nucleotide changes. Genomic positions of mutations were determined relative to the amplicon start position. We then compared observed mutations to predicted editing sites derived from prior computational analyses of sgRNA targets. To assess consistency across experimental replicates, we examined sequence intersections and calculated allele frequencies at two timepoints (Day 5 and Day 17).

### Library Sequencing Analysis

Guide sequences were extracted from the FASTA file of the library sequencing. First, the U6 promoter and gRNA scaffold TTGTGGAAAGGACGAAACACC…GTTTCAGAGCTATGCT was trimmed using cutadapt. High-quality trimmed reads were then aligned against the FASTA file of all sgRNA sequences using bowtie ^45^. The aligned reads were processed using custom code (available on GitHub) to obtain the read counts of sgRNAs.

Before proceeding to correlation coefficient calculation, read counts were first normalized to log2reads per million using

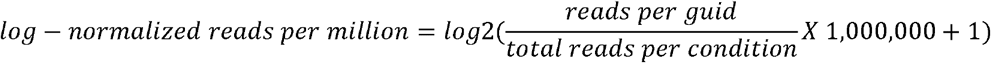

PyDESeq2 was installed in a separate conda environment following the instructions: https://pydeseq2.readthedocs.io/en/latest/. DESeq2 uses the Benjamini-Hochberg method for multiple hypothesis testing. We then used *PyDESeq2* to determine sgRNA log2foldchange depletion. All dropout comparisons were made to the D0 baseline library.

### sgRNA editing efficiency estimation

We utilized the *BE-Hive* package^46^ to predict the editing efficiencies of sgRNAs in mouse embryonic stem (mES) cells, the only suspension cell line in their dataset that closely resembles K562 and Jurkat cells. ABE sgRNA efficiencies were predicted using the ABE model, while CBE sgRNA efficiencies were predicted using the BE4 model. The BE-Hive tool generates a logit score for each sgRNA, representing its propensity for base editing. This score is then transformed into a fraction—corresponding to the proportion of sequenced reads exhibiting base editing activity—using a logistic function.

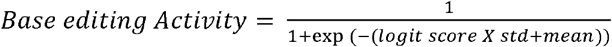. According to the iSilence control, we use an average editing efficiency of 30% to perform the transformation (default std = 2). For each sgRNA, we calculated the adjusted fraction of edited reads based on this efficiency. sgRNAs with reading counts falling below two standard deviations from the mean were excluded from further analysis.

### Estimation of fold Difference and Likelihood Ratio Test

To quantify the relative probability of fitness effects between synonymous and non-synonymous sgRNAs, we modeled the observed hit rates in our pooled screen using a binomial framework. The maximum likelihood estimates (MLE) for the probability of a fitness effect are 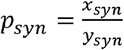 and 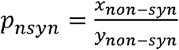 synonymous mutations under binomial model, where X represents the number of validated hits and Y the estimated number of screened sites for each mutation category. To estimate the fold difference between synonymous and missense variants, we calculated the likelihood ratio (r) as the ratio of the observed proportions. Given the extremely low event count in the synonymous group, we chose to maximize the joint likelihood for each r to calculate the confidence interval for r. The likelihood ratio test statistics was as calculated for each r as *LRT(r) = 2*[*logL*(*p_syn_, p_non-syn_*) − *logL*(*r*)]. The confidence interval was then defined as 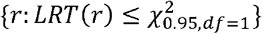.

To statistically test the significance of the likelihood ratio, we conducted a likelihood ratio test. The parameters *P_syn_*and *P_non-syn_*are treated as hyperparameters across the plotted surface. For each grid point (*p_syn_, p_non-syn_*) we computed the joint log-likelihood of the observed data under the assumption that synonymous and non-synonymous mutations have different probabilities of being functional by

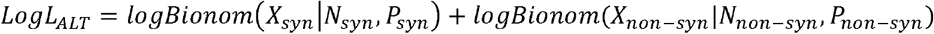

We then compared this alternative model to the null model, which assumes a shared probability of fitness effect for both mutation types, 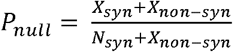 by computing the likelihood ratio test statistics *LRT = 2[logL_Alt_*(*P_syn_, P_non-syn_*)*− logL_null_]*. P-values were derived from the *χ*^2^, df = 1 distribution.

## AUTHOR CONTRIBUTIONS

**Y.R**: Conceptualization, investigation, methodology, data analysis, writing and editing. **I.S**: Methodology. **J.R.P**: Conceptualization, supervision, funding-acquisition, project administration, writing and editing. **E.P.O**: Conceptualization, supervision, funding-acquisition, project administration, writing and editing.

## COMEPETING INTERESTS

JRP is a co-founder of RedAce Bio. JRP was a co-founder and consultant for Theseus Pharmaceuticals. JRP held equity in Theseus Pharmaceuticals. JRP holds equity in MOMA therapeutics and RedAce Bio. JRP has consulted/consults for MOMA therapeutics, Curie.Bio, Third Rock Ventures, Takeda Pharmaceuticals, Galapagos Pharmaceuticals, and Roche/Genentech. JRP has received honoraria and travel expenses from Roche/Genentech, Third Rock Ventures, and Theseus Pharmaceuticals.

## DATA AND MATERIEALS AVAILABILITY

The base editor library and single sgRNA sequencing data are deposited under Bioproject PRJNA1236376. All custom code used for analysis is available on GitHub (https://github.com/ryy1221/synSg).

